# SCANNER: A Web Resource for Annotation, Visualization and Sharing of Single Cell RNA-seq Data

**DOI:** 10.1101/2020.01.25.919712

**Authors:** Guoshuai Cai, Feifei Xiao

## Abstract

**Motivation:** In recent years, efficient scRNA-seq methods have been developed, enabling the transcriptome profiling of single cells massively in parallel. Meanwhile, its high dimensionality brought challenges in data modeling, analysis, visualization and interpretation. Available analysis tools require extensive knowledge and training of data properties, statistical modeling and computational skills. It is challenging for biologists to efficiently view, browse and interpret the data.

**Results:** Here we developed SCANNER, as a public webserver resource to equip the biologists and bioinformatician to share and analyze scRNA-seq data in a comprehensive and collaborative manner. It is effort-less and host-free without requirement on software setup or coding skills, and enables a user-friendly way to compare the activation status of gene sets on single cell basis. Also, it is equipped with multiple data interfaces for easy data sharing and currently provide a database for studying the smoking effect on single cell gene expression in lung. Using SCANNER, we have identified larger proportions of cancer-associated fibroblasts cells and activeness of fibroblast growth related genes in melanoma tissues in females compared to males. Moreover, we found *ACE2* is mainly expressed in pneumocytes, secretory cells and ciliated cells with disparity in gene expression by smoking behavior.

**Availability and implementation:** SCANNER is available at https://www.thecailab.com/scanner/.

**Supplementary information:** Supplementary data are available online.

**Key Points:**

- SCANNER provides a new web server resource for promoting scRNA-seq data analysis
- SCANNER enables comprehensive and dynamic analysis and visualization, novel functional annotation and activeness inference, online databases and easy data sharing.
- SCANNER bridges the data analysis and the biological experiment units.

## Introduction

In recent years, single cell RNA sequencing (scRNA-seq) methods have been efficiently enabling the transcriptome profiling of each single cell (Kolodziejczyk, et al., 2015). It provides opportunities to identify cell clusters and their specific biomarkers and insights into cell developmental trajectory, gene bursting activities, cell interactions and others (Hwang, et al., 2018). However, the high dimensionality of scRNA-seq data also brought challenges in its analysis. A widely used strategy is to reduce the high dimension into a low space of two- or three-dimensions, and in which, data analysis, visualization and interpretation could be further performed. For fast browsing scRNA-seq data, applications such as SCV (Wang, et al., 2019) and Cerebro (Hillje, et al., 2019) have been developed. However, they require local installation and implementation, as well as a multi-step pre-processing procedure including raw data processing, dimension reduction and data input standardization. This brings obstacles for biologists to easily explore the data and efficiently communicate with bioinformatician whose knowledge of data modeling and computational skills is much needed. Moreover, due to the rapid development of new technologies, the volume and complexity of data increase fast and highly require an effective gateway for data sharing, storage, annotation and visualization. The Broad Institute Single Cell Portal (https://portals.broadinstitute.org/single_cell) and scRNASeqDB (Cao, et al., 2017) are available databases for scRNA-seq studies. However, their implemented functions are static and limited for data visualization and analysis. A new web server resource with comprehensive and dynamic analysis and visualization functions is highly demanded to bridge the data analysis and the biological experiment units.

## Features

In this study, we developed the <u>S</u>ingle <u>C</u>ell Transcriptomics <u>Ann</u>otated View<u>er</u> (SCANNER), as a public web resource for scRNA-seq data management, analysis and interpretation in a comprehensive, flexible and collaborative manner. SCANNER is based on the framework of SCV, which is a useful local R Shiny application for scRNA-seq data visualization. SCANNER enables a unique set of functions highlighted in Figure 1, adding complimentary value to SCV and other existing tools. Specifically,

1. SCANNER is a web server application. Compared to SCV and other local applications, it requires no software setup and enables host-free work, fulfilling the demand of efficient communication for seamless collaboration. Compared to existing web applications, SCANNER enables a flexible and dynamic comparison between groups, comprehensively with five visualization modules for identified clusters (Cluster), gene expression in cells (Expression), distribution of gene expression in clusters (Distribution), expression detection rate and size in clusters (Detection) and expression heatmap with hierarchical clustering across genes and cells (Similarity). Each module dynamically allows parameters setting for object selection, group comparison, cluster filtering and layout setting. To maintain a favorable computational speed and resource usage, SCANNER currently visualizes a maximum of 500 cells per cell cluster for large datasets (Supplementary Methods), which should effectively represent the cell population in a cluster.
2. SCANNER provides novel functional annotation and activeness inference. Analysis on gene set can provide valuable insights into explaining biological mechanisms underlying a phenotype of interest. To enable this, SCANNER infers the activation status of a particular gene set that are involved in a pathway by four scores (Supplementary Methods), including averaging expression abundance or ranks of the involved genes, the expression of an eigengene (Langfelder and Horvath, 2008), and enrichment scores (Barbie, et al., 2009).
3. SCANNER provides an online database for scRNA-seq data. Currently, SCANNER hosts a database of smoking lungs for studying the smoking effect on single cell basis. Studies has been reviewed and three publicly available datasets were included, including the dataset from bronchial epithelial cells, ALCAM+ epithelial cells and CD45+ white blood cells from six never and six current smokers (Duclos, et al., 2019), the dataset from lung tissue of one former smoker, two current smoker and five non-smokers) (Reyfman, et al., 2019) and the dataset from lung tissue of one former smoker, one current smoker and three non-smokers (Madissoon, et al., 2019). We will continue to collect datasets and update SCANNER to provide a comprehensive resource of single cell transcriptomics for public use.
4. SCANNER provides multiple data interfaces which enables easy data sharing. Three options are available, by (i) SCANNER data object that users can generate following instruction (Availability and implementation), (ii) access credentials that users can contact the authors for a password controlled access; and (iii) database that provides the most efficient method for data sharing and exploring.

**Figure 1.**
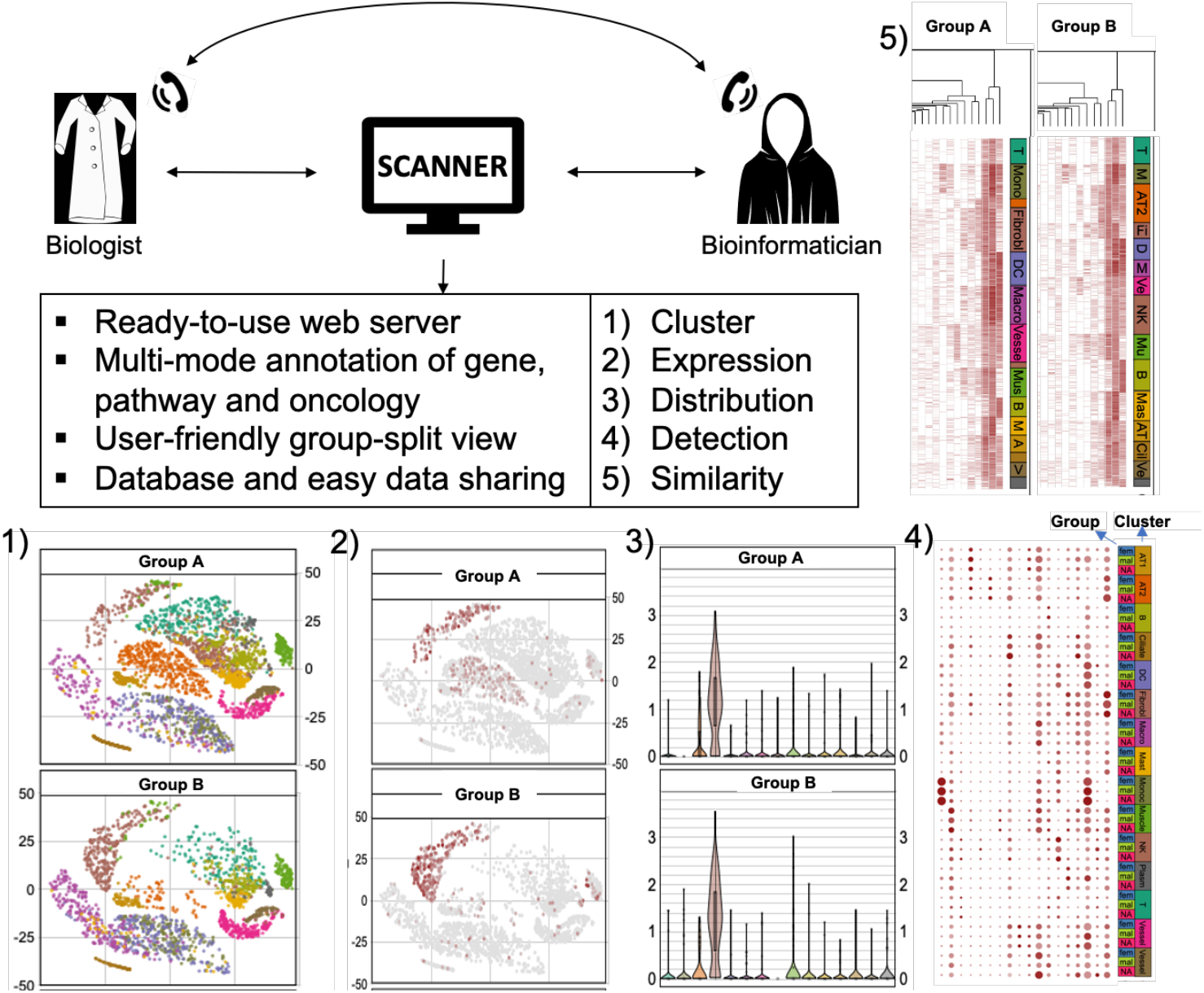
SCANNER application scenario. Scanner facilitates the sharing, visualization, analysis and interpretation of scRNA-seq data and the communication between biologists and bioinformatician in a flexible, multi-functional and user-friendly manner. Its group side-by-side view is useful to explore differences between groups, of 1) cell clusters, 2) single cell gene expression or pathway activity, 3) expression/activity distribution, 4) expression/activity detection and 5) expression pattern.

## Case studies

Here, we demonstrate the application of SCANNER with two case studies:

a. **Sex disparity in melanoma-associated fibroblast.** We analyzed the melanoma dataset (Tirosh, et al., 2016) and found that tumors from female patients had significantly larger proportions of cancer-associated fibroblasts (CAF) and endothelial cells than those from males (Fig. S1A). Correspondingly, in CAF and endothelial cells of the female patients, the fibroblast growth factor (FGF) binding function inferred by enrichment score was highly activated (Fig. S1B, C) with higher detection rates and activeness scores (Fig. S1D), which were confirmed by methods of average expression, average rank and eigen-gene expression (Fig. S2). Consistently, the FGF genes FGF1 and FGF2 were highly expressed in CAF cells in the female melanoma tissue (Fig. S3). Such over-expression was also found in most of genes involved in the FGF binding function (Fig. S4, 5). Given that CAF is a promise target to treat melanoma (Zhou, et al., 2015), this gender difference in CAF may provide a new implication to explain why male patients usually have worse survival outcome compared to females (Joosse, et al., 2013).
b. **Tobacco-use disparity in *ACE2* lung expression.** Exploring our database of smoking lung, we found that the gene of the SARS-CoV-2 receptor, ACE2, is mainly expressed in pneumocytes, secretory cells and ciliated cells (Fig. S6), which is consistent with the recent study of Ziegler et al. (Ziegler, et al., 2020). Also, among bronchial epithelial cells, we found that ACE2 gene is mainly expressed in club cells in never smokers. Differently in smokers, goblet cells are extensively proliferated and harbor most expressed ACE2, which may indicate a complex effect of smoking on the COVID-19 risk (Cai, et al., 2020).

## Acknowledge

We acknowledge Ben Torkian and Jun Zhou from Research Computing program of University of South Carolina for the assistance on gateway application, allocation and implementation. This study was supported by the NSF XSEDE Startup Allocation Award (MCB190139).

