## Supplementary information for "SCANNER: A Web Resource for Annotation, Visualization and Sharing of Single Cell RNA-seq Data"

**Short title:** A Web Server for scRNA-seq Data

**Corresponding Authors:**

Guoshuai Cai, PhD
Department of Environmental Health Sciences
Arnold School of Public Health
University of South Carolina
921 Assembly Street
Public Health Research Center 401C
Columbia, SC  29204


Feifei Xiao, PhD
Department of Epidemiology and Biostatistics
Arnold School of Public Health
University of South Carolina
915 Greene Street, Room 449
Columbia, SC  29204


### Supplementary Methods

Gene set activity inference

Four approaches are available in SCANNER for inferring the relative activeness of a particular pathway, function or set of interested genes across cells. In one of all $q$ cells, the activeness score $S$ of an interested gene set, which has $m$ genes for the $i$-th gene, is calculated by,

- *Average expression*, that

$$S = \frac{1}{m}\sum_{i=1}^{m} E_i$$

where $E_i$ is the scRNA-seq normalized data of the $i$-th gene. We used Seurat normalized sequencing count in *log* scale in developing the smoking lung database.

- *Average rank, that*

$$S = \frac{1}{m}\sum_{i=1}^{m} R_i$$

where $R_i$ is the rank of the $i$-th gene.

- *Eigen-gene expression,* which can be considered as a weighted average expression from a gene set. Briefly, the covariance matrix $C_{m\times m}$ of the gene set expression matrix is calculated by

$$C_{m\times m} = \frac{1}{q-1}X^T X$$

where $X$ is the zero centered and unit variance scaled expression matrix of $q \times m$ size, where $q$ is the number of cells and $m$ is the number of genes of interest. Since the covariance matrix $C$ is symmetric and positive definite, the eigenvalues of $C$ are real and it can be diagonalized as

$$C = V\Lambda V^T$$

where $V$ is the eigenvector matrix and $\Lambda$ is an eigenvalue matrix. The principle components (PCs) can be obtained by $XV$ and $S$ is given by the first PC (PC1) which represents the most variation in the data. To reduce the effect of extreme values on SCANNER visualization, we constrain $S$ within its $5^{th}$-$95^{th}$ percentile.

- *Gene set enrichment score,* which is calculated using a similar strategy of single sample gene set enrichment analysis (ssGSEA) (Barbie, et al., 2009). For the gene set $G$ of size $m$ from all $n$ genes in a sample,

$$S = \sum_{j=1}^{n}\left[\sum_{i\in G, i\leq j}\frac{{E_i}^{\alpha}}{\sum_{i\in G}{E_i}^{\alpha}} - \sum_{i\notin G, i\leq j}\frac{1}{(n-m)}\right],$$

where $E_i$ gives more weight on highly expressed genes, and $\alpha$ controls the degree of the weight. In this study, SCANNER set $\alpha = 1$ which was typically used in the regular ssGSEA.

Current activity inference is available for MSigDB v.7.0 (Liberzon, et al., 2011) gene sets of Hallmark Collection, KEGG Pathway, Biocarta Pathway, Reacome Pathway, GO Biological Process, GO Cellular Component, GO Molecular Function, Oncogenic Signatures and Immunological Signatures.

Cell subsetting

For large-scaled datasets, SCANNER subset cells to maintain a favorable computing speed and resource usage. Currently, maximum 500 cells for each cluster are selected by SCANNER using two approaches:

*Random sampling*: cells in each cluster are randomly selected.

*Prediction ellipse*: cells in an fitted ellipse which surround a particular cluster at a quantile of max(1, $\frac{500}{the\ cell\ number\ in\ cluster}$) are selected. The ellipse can be easily defined by its covariance matrix and vector of means in a low-dimensional space of scRNA-seq data. SCANNER used -distributed Stochastic Neighbor Embedding (t-SNE) (van der Maaten and Hinton, 2008) dimensions in developing the smoking lung database. The axis scales of ellipse were calculated using the square root of the eigenvalues of the covariance matrix and then the distance to ellipse can be solved. R Package "SIBER" was used for realizing this function. This method selects the core cells that are most distinct for each cluster.

### Supplementary Figures

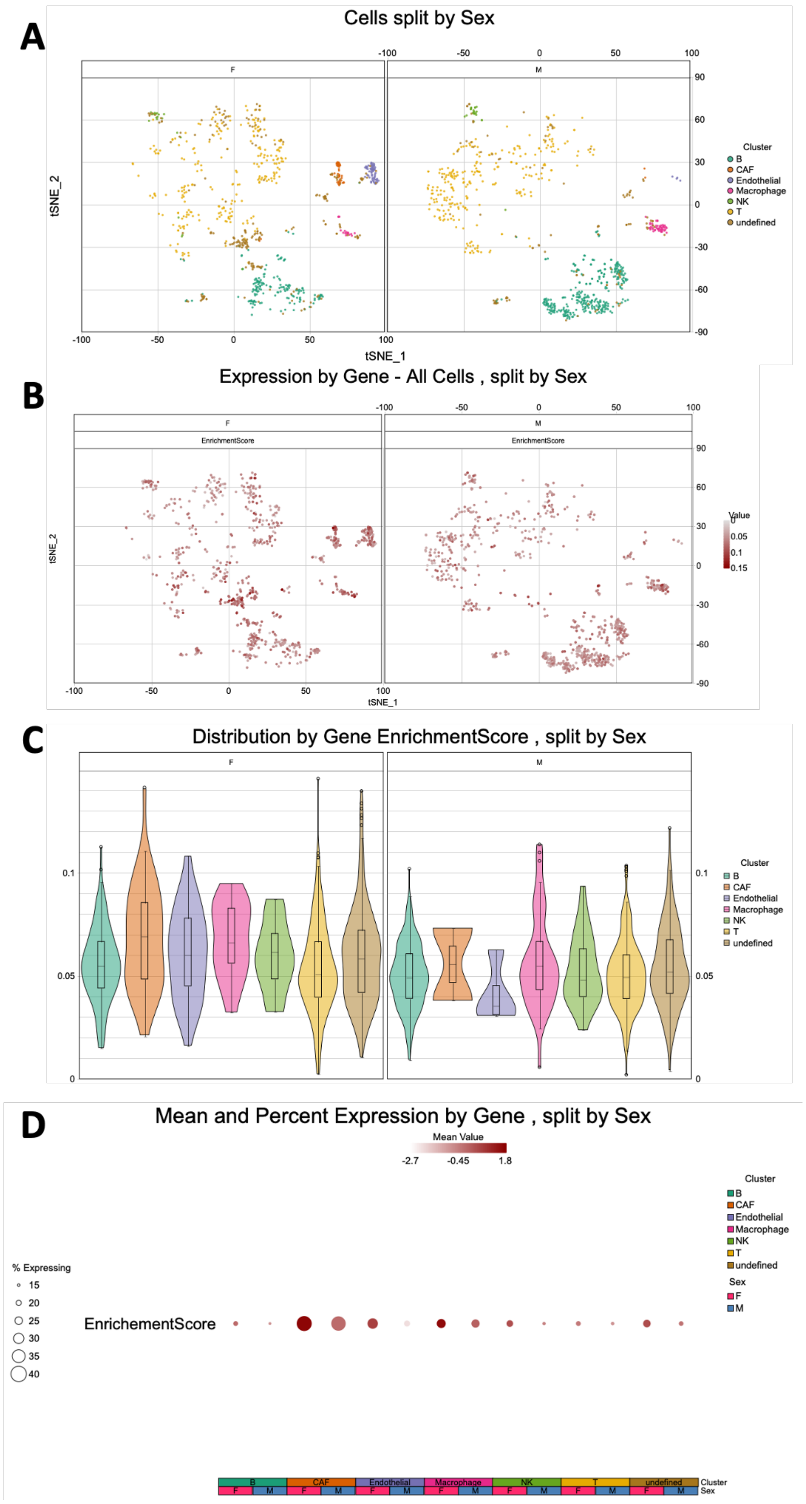


**Figure S1. SCANNER identify sex disparity in melanoma-associated fibroblast.** The disparity is found in (A) cell cluster, (B) enrichment score of GO: fibroblast growth factor binding function, (C) enrichment score distribution and (D) enrichment score level and detection rate. Darker color indicates higher activity in (B) and (D) and a larger size indicates a higher detection rate in (D). F=Female, M=Male.

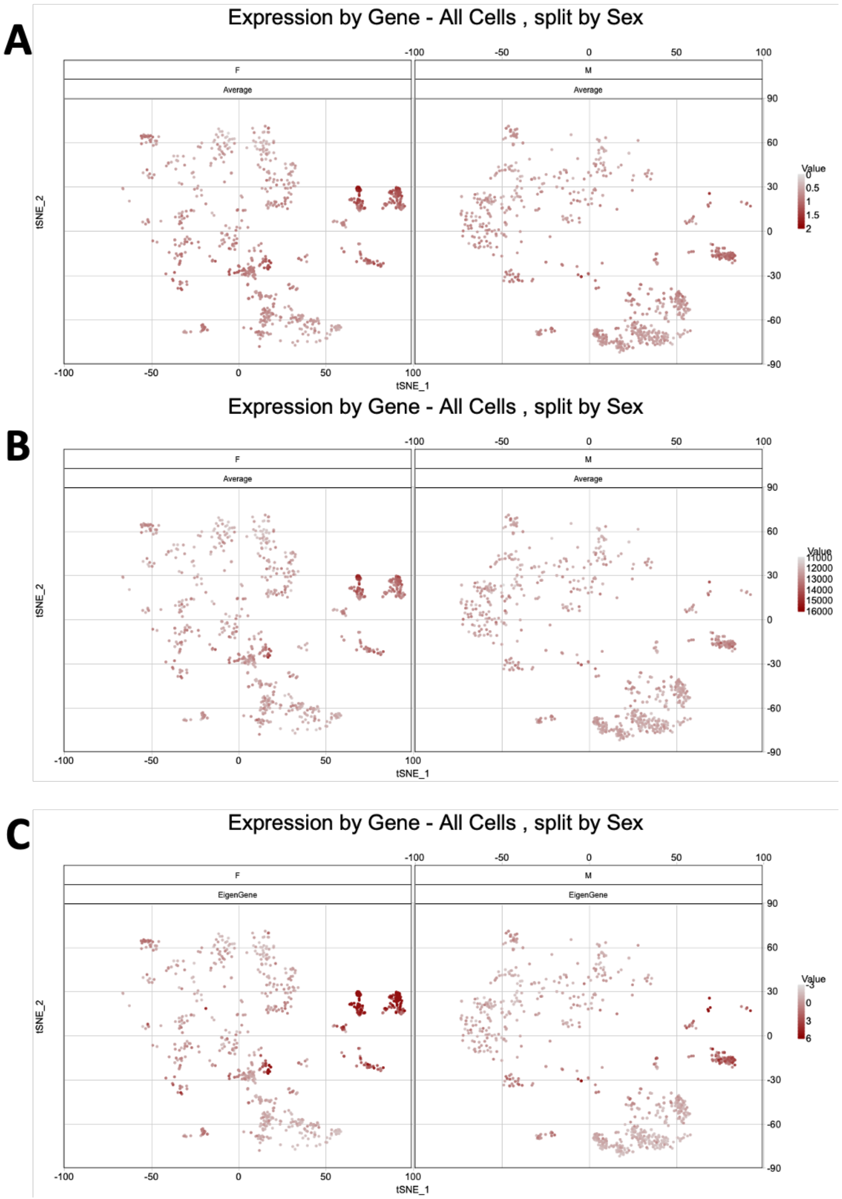


**Figure S2. Sex disparity in activeness of fibroblast growth factor binding.** The disparity is found in GO: fibroblast growth factor binding function activeness inferred by (A) average expression, (B) average rank and (C) eigen-gene expression. Darker color indicates higher activity. The cell clusters with darker color in the F group are cancer-associated fibroblasts and Endothelial cells. F=Female, M=Male

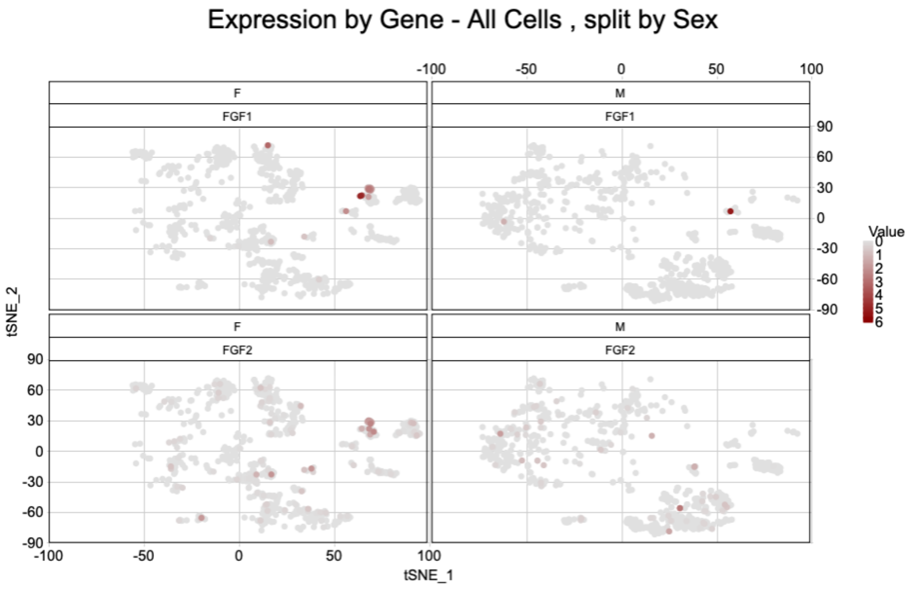


**Figure S3. Sex disparity in expression of *FGF1* and *FGF2* expression.** Darker color indicates higher expression. The cell clusters with darker color in the F group are cancer-associated fibroblasts and Endothelial cells. F=Female, M=Male.

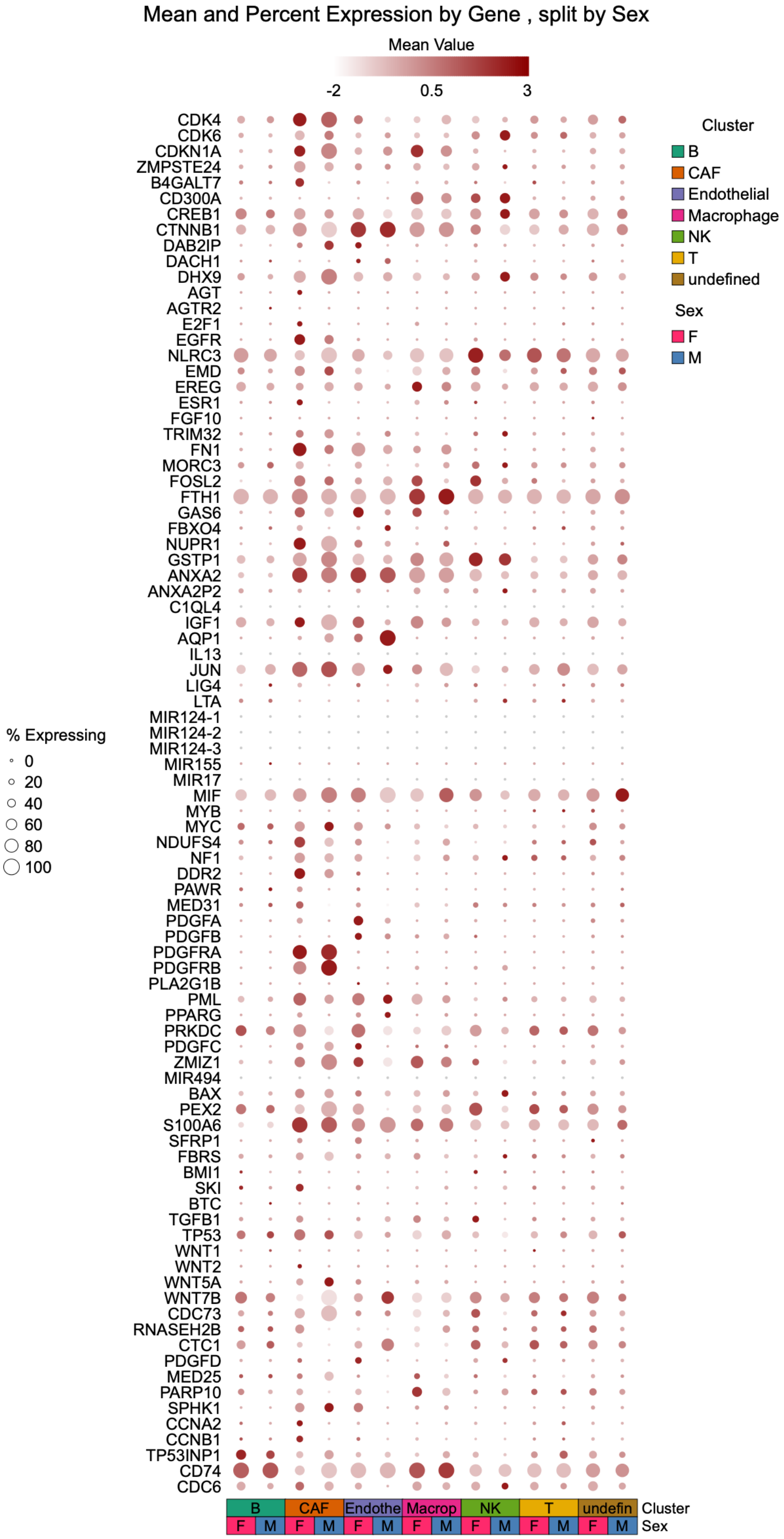


**Figure S4. Sex disparity in expression of fibroblast growth factor binding related genes.** Darker color indicates higher expression. Larger size indicates higher detection rate. F=Female, M=Male.

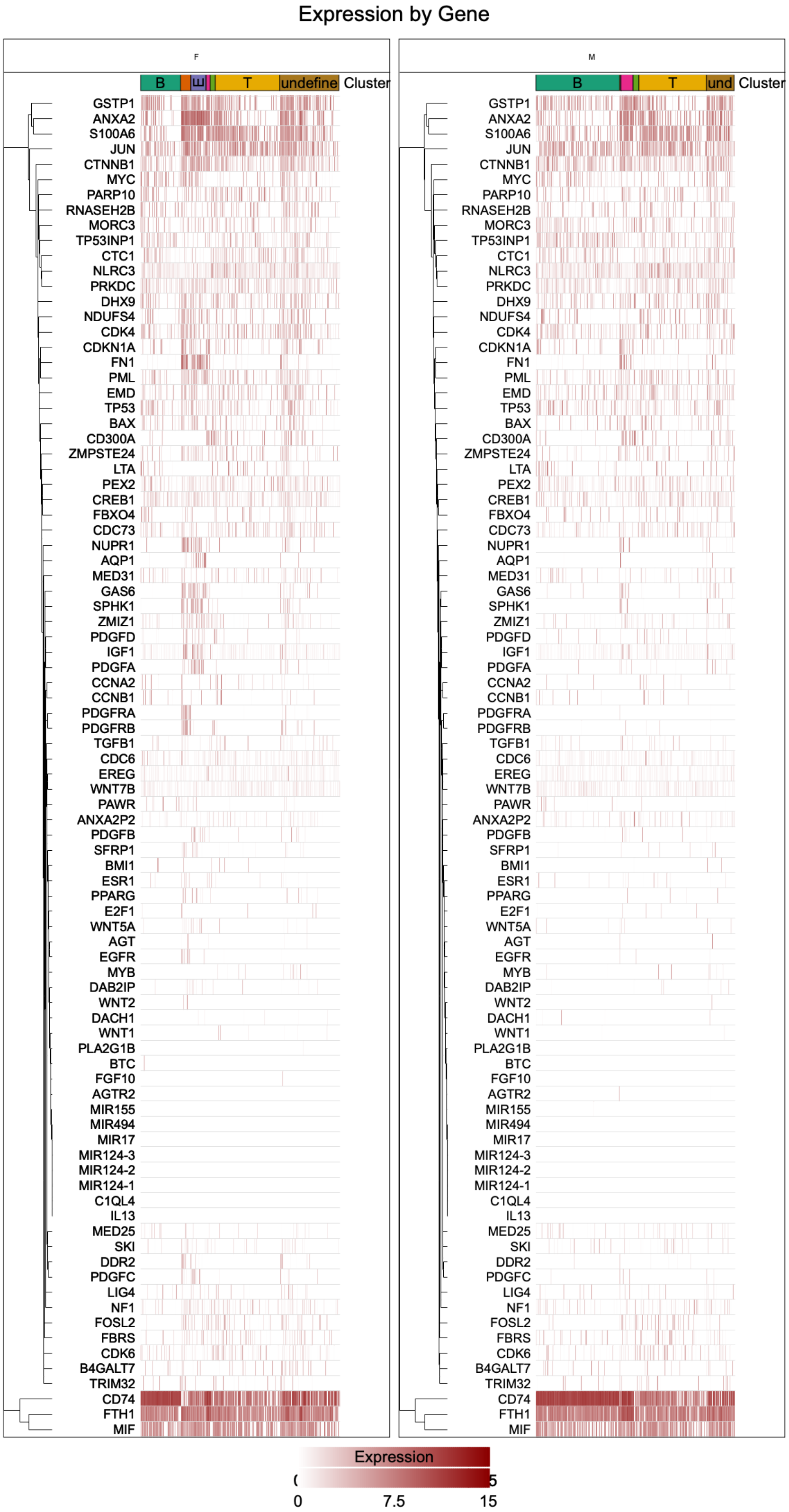


**Figure S5. Sex disparity in expression pattern of fibroblast growth factor binding related genes.** Darker color indicates higher expression. F: Female, M: Male.

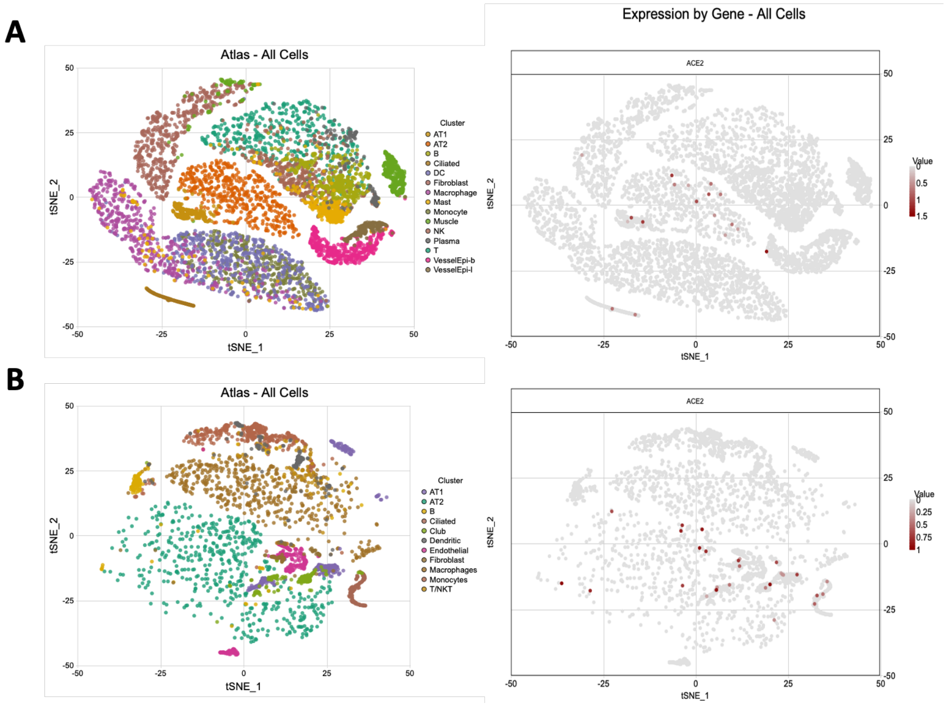


**Figure S6. SCANNER detected *ACE2* expression in specific lung cells.** Two datasets including (A) the GSE122960 dataset and (B) the Meyer dataset shows *ACE* expression (right panel) mainly expressed in pneumocytes, secretory cells and ciliated cells (left panel).
